# Protease-controlled secretion and display of intercellular signals

**DOI:** 10.1101/2021.10.18.464444

**Authors:** Alexander E. Vlahos, Jeewoo Kang, Carlos A. Aldrete, Ronghui Zhu, Lucy S. Chong, Michael B. Elowitz, Xiaojing J. Gao

## Abstract

To program intercellular communication for biomedicine, it is crucial to regulate the secretion and surface display of signaling proteins. If such regulations are at the protein level, there are additional advantages, including compact delivery and direct interactions with endogenous signalling pathways. We created a modular, generalizable design called <u>**R**</u>etained <u>**E**</u>ndoplasmic C<u>**lea**</u>vable <u>**Se**</u>cretion (RELEASE), with engineered proteins retained in the endoplasmic reticulum and displayed/secreted in response to specific proteases. The design allows functional regulation of multiple synthetic and natural proteins by synthetic protease circuits to realize diverse signal processing capabilities, including logic operation and threshold tuning. By linking RELEASE to additional novel sensing and processing circuits, we were able to achieve elevated protein secretion in response to “undruggable” oncogene KRAS mutants. RELEASE should enable the local, programmable delivery of intercellular cues for a broad variety of fields such as neurobiology, cancer immunotherapy and cell transplantation.

## Introduction

Synthetic biology aspires to create biomolecular circuits that can sense the state of cells, process the information, and then deliver therapeutic outputs accordingly^1,2^ This vision has been enhanced by the creation of protein-based circuits by others^3–6^ and ourselves^7^. Protein-based circuits have advantages such as fast operation, compact delivery, and robust, context-independent performance compared to traditional transcriptional circuits^6,7^. However, these protein circuits have operated in the cytosol, and there remains an urgent need for a design that enables protein-level control of intercellular communication, often required at the “respond” step in “sense-process-respond”.

Cell-cell communication is widespread^8–10^ and essential for diverse biological processes, such as the generation of immunological responses^11,12^, cell differentiation and tissue development^13–15^, the maintenance of physiological homeostasis^16^, and cancer development^17,18^. Intercellular communication is typically implemented by secreted molecules, including hormones and cytokines. To take cancer immunotherapy as an example, there an ideal application would be introducing a protein circuit that sense the cancerous state of a cell, secrete immunostimulatory signals with temporal and quantitative precision to mobilize the immune system while lysing the cell, and therefore turn these cells into vaccines against other similarly cancerous cells. This would not only avoid the toxic effects associated with the systemic delivery of immunomodulating proteins^19,20^, but also match the complex, dynamic immune process we are trying to control^21^. In contrast, of the current local delivery methods^22^, neither nanoparticle^23^, or biomaterial-based^24^ delivery platforms can fulfill the aforementioned functions that circuits can deliver.

Given the importance of intercellular communication, we sought to interface protein circuits with the secretion and display of protein signals. Specifically, because protease activity has emerged as a “common currency” of protein circuits that responds to synthetic and endogenous inputs^6,7,25–28^, it will be ideal to directly control protein secretion using proteases. To design a modular protease-regulated protein secretion system, we adapted aspects of the natural secretion process. Secreted proteins are typically transported into the Endoplasmic Reticulum (ER), processed in the Golgi apparatus, and finally secreted at the plasma membrane. Some proteins contain signaling motifs (e.g., KDEL for soluble proteins^29^ and the cytosol-facing dilysine (-KKXX) or -RXR motifs for membrane proteins^30–34^) recognized in the early Golgi apparatus, causing the protein to be retrieved, transported retrogradely, and retained in the ER^32,35^. Other ER-resident proteins, such as cytochrome p450 are retained in at the ER via their signal-anchor transfer sequence^36,37^. These retention motifs function in their endogenous contexts as well as when fused to normally secreted proteins^30,33^. To place ER retention under protease control, we engineered the modular <u>**R**</u>etained <u>**E**</u>ndoplasmic C<u>**lea**</u>vable Secretion (RELEASE) platform, compatible with both protein secretion and the surface display of membrane proteins. We validated and optimized the core mechanism of RELEASE, created input-processing capabilities, and then used RELEASE to control physiological outputs. Finally, we combined RELEASE with novel sensing and processing components to respond to internal cell states and external signals via engineered receptors. This study demonstrates a novel protein-level control module to directly regulate protein secretion that is compatible with pre-existing protein components to program therapeutic circuits for cancer immunotherapy and transplantation in the future.

## Results

### Engineering RELEASE for protein secretion and expression

RELEASE contains 4 components: a luminal facing linker containing a furin endoprotease cut site, a transmembrane anchor domain^38^, a cytosolic linker containing a protease cleavage site, and an ER retention motif (**Fig. 1a, 1b**). On the cytosolic face, the retention motif ensures that the tagged protein is actively transported back to ER^39,40^, a process only aborted after the motif is removed by a proteases such as tobacco etch virus protease (TEVP)^6,7^. On the luminal face, soluble proteins are initially tethered to the membrane through the linker and thus coupled to the cytosolic ER retention signal^7^. After the first cytosolic cleavage event, the membrane-tethered protein is processed into its soluble form through cleavage by furin in the trans-Golgi apparatus (furin is absent in cis-Golgi or ER)^41^ (**Fig. 1a**), and finally secreted.

**Figure 1:**
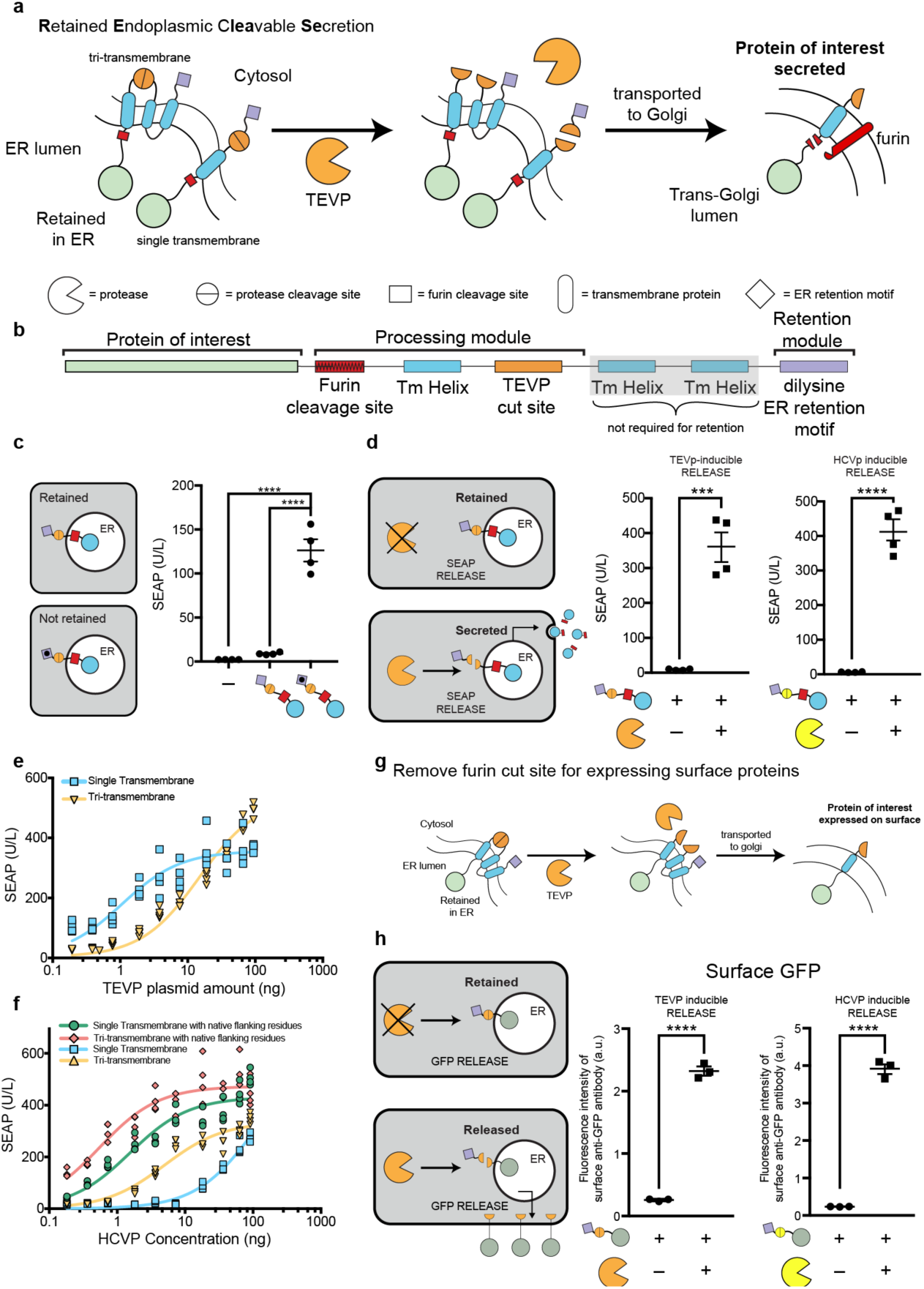
Design of Retained Endoplasmic Retained Secretion (RELEASE). **a)** Proteins of interest are fused to RELEASE and retained in the ER via the dilysine ER retention domain (purple diamond). Upon activation or expression of a protease such as TEVP (orange partial circle), the ER retention domain is removed (middle panel) and the protein of interest is transported through the constitutive secretory pathway. When reaching the Trans-Golgi Apparatus (right panel), the native furin endoprotease cleaves the linker region allowing the membrane-bound protein to be secreted. **b)** RELEASE is a modular platform and can be modified to respond to different proteases and regulate different proteins of interest. **c)** The C-terminal dilysine motif of RELEASE is required for SEAP retention and mutation of the two lysine residues to alanines (KKXX-COOH → AAXX-COOH) increased SEAP secretion. There was no significant difference in signal between RELEASE and control cells without SEAP. **d)** Co-expression of proteases such as TEVP (orange partial circle), or HCVP (yellow partial circle) with the respective RELEASE constructs increased SEAP secretion. **e)** Single transmembrane and tri-transmembrane RELEASE constructs had different cleavage efficiencies to TEVP cleavage. **f)** The cleavage efficiencies of HCVP RELEASE constructs were also affected by transmembrane selection and was improved by modifying the residues flanking the HCVP cut site with native linker proteins. Based on the steady-state solution of a kinetic model for proteolytic cleavage, we determined that the relation between RELEASE output and the amount of protease plasmids fits the Michaelis-Menten equation^7^. We therefore fit the titration curves using Michaelis-Menten equations and used K_m_ to represent the apparent cleavage efficiency of each design by its corresponding protease. A complete list of the calculated cleavage efficiencies for the different RELEASE constructs can be found in **Supplementary Table 1. g)** By removing the furin cut site, RELEASE was amenable to control the surface display of proteins. **h)** Increased surface display of membrane-bound GFP fused to RELEASE in response to TEVP (left panel) or HCVP (right panel). Each dot represents a biological replicate. Mean values were calculated from four (**c-f**) or three replicates (**h**). The error bars represent +/- SEM. The results are representative of at least two independent experiments; significance was tested using an unpaired two-tailed Student’s *t*-test between the two indicated conditions for each experiment. For experiments with multiple conditions, a one-way ANOVA with a Tukey’s post-hoc comparison test was used to assess significance. *** = p < 0.001, **** = p < 0.0001

First, to validate the effectiveness of the retention motif, we fused it to secreted embryonic alkaline phosphatase (SEAP)^42^, and used a dilysine-lacking mutant motif as the negative control. We transfected human embryonic kidney (HEK) 293 cells using DNA plasmids encoding the constructs. Using RELEASE, SEAP is minimally present in the supernatant and comparable to control cells that were not transfected with SEAP (**Fig. 1c**). Mutation of the dilysine motif of RELEASE significantly increases SEAP secretion (**Fig. 1c**). We next placed the dilysine motif under the control of TEVP, and showed that the co-expression of TEVP significantly increases SEAP secretion (**Fig. 1d – left panel**). By switching the cytosolic protease cut sites, we validated RELEASE against other orthogonal proteases such as the hepatitis C virus protease (HCVP) (**Fig. 1d – right panel**) and the tobacco mottling vein virus protease (TVMVP) (**Supplementary Fig. 1a b**). Furthermore, the design is compatible with alternative ER-retention motifs, as we validated constructs using the N-terminal signal anchor sequence from p450^36,37^ (**Supplementary Figure 2a b**).

In anticipation of tuning RELEASE for different applications, we next explored how its performance is affected by two design decisions. First, as an alternative to the tri-transmembrane domain^38^, we created a single transmembrane variant, and found it more sensitive to TEVP compared to the tri-transmembrane construct (**Fig. 1e**). Similarly, the input sensitivity of HCVP-inducible RELEASE is also modulated by the choice of the transmembrane domain (**Fig. 1f**). Furthermore, by using a protein linker containing the native residues that flank the HCVP cut site^38,43^, we made more sensitive HCVP-inducible RELEASE constructs (**Fig. 1f – red and green lines**) than the original versions that use synthetic flanking sequences. A complete list of the cleavage efficiencies for the RELEASE variants are in **Supplementary Table 1**. We took advantage of this tunability to reduce RELEASE response to the input-independent activity of a membrane-localized split protease^44^ (**Supplementary Fig. 3a**) and therefore improve output dynamic range (**Supplementary Fig. 3b, c**).

In addition to controlling protein secretion, cells can communicate by changing the display of proteins on their surface^12,13^ By removing the furin cut site in RELEASE, we hypothesized it could control the surface display of proteins (**Fig. 1g**). To validate this strategy, membrane-bound green fluorescent protein (GFP) fused to RELEASE was transfected into HEK293 cells, and the cell surface was stained using an anti-GFP antibody. GFP-RELEASE constructs significantly increased surface display of GFP in response to the cognate proteases (**Fig. 1h**). Taken together, these results show that RELEASE is a suitable approach to control the secretion and surface display of proteins in response to protease activity (**Fig. 1d, h)**.

### RELEASE is compatible with circuit-level functions

After validating the RELEASE design, our next goal was to ensure that its activation could be programmed using protease-based circuits, either pre-existing^6,7^ or novel. For RELEASE to operate properly in circuits with multiple proteases, first it is important to validate the orthogonal control of RELEASE by the selected protease^7^. Indeed, cells simultaneously transfected with two RELEASE constructs (**Fig. 2a**) were orthogonal and only secreted the respective reporter protein in response to the cognate protease (**Fig. 2b**). This result demonstrates that two proteases can be used to independently regulate secretion of distinct target proteins in the same cell.

**Figure 2:**
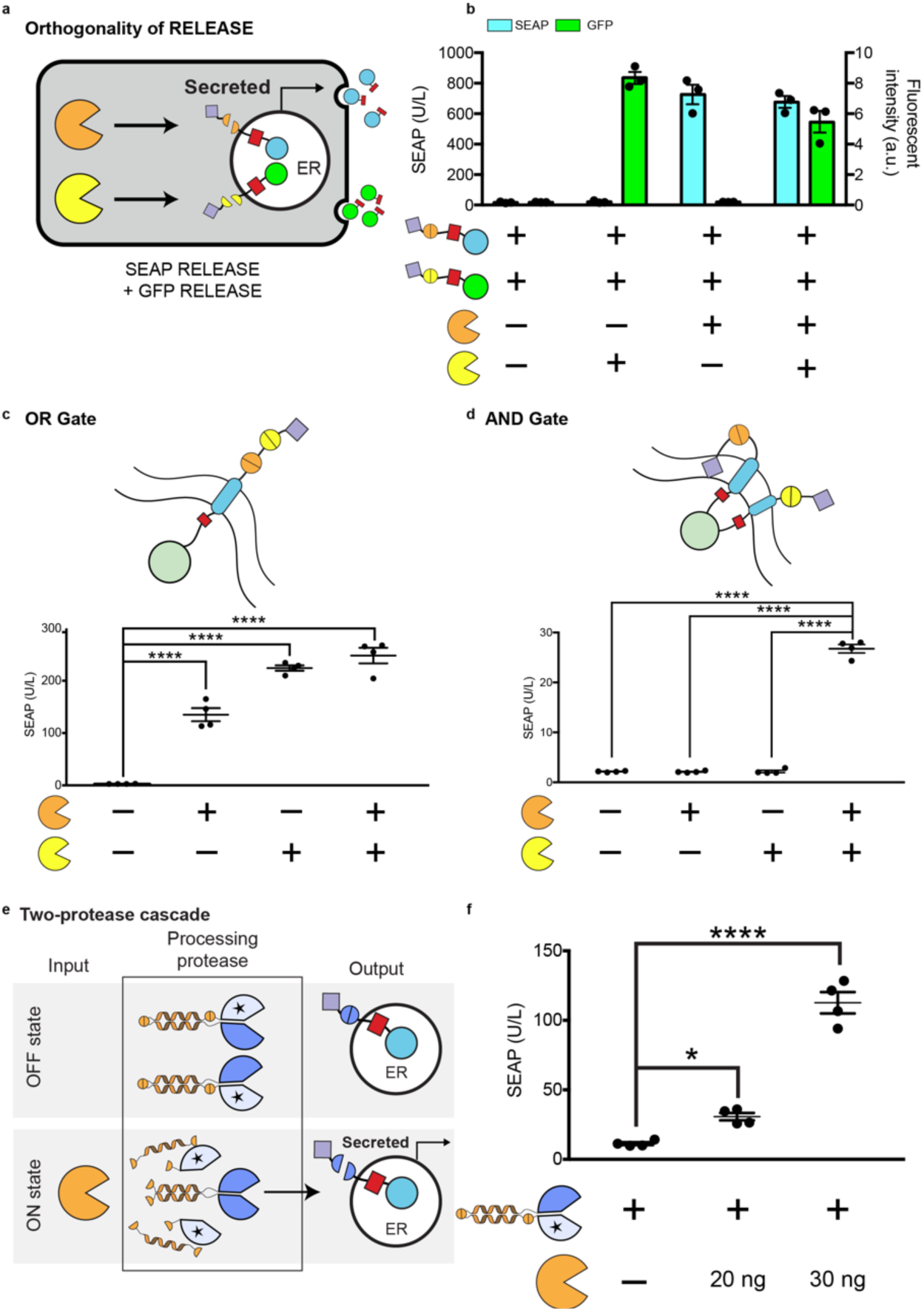
RELEASE in circuits. **a)** Orthogonal operation of RELEASE constructs. **b)** HEK293 cells were co-transfected with SEAP fused to RELEASE (responsive to TEVP) and GFP fused to RELEASE (responsive to HCVP). SEAP and GFP levels increase in the supernatant when the cognate protease was expressed. **c)** Tandem insertion of two protease cut sites (top panel) created a RELEASE construct that implemented OR gate logic. If either of the respective proteases were expressed, the dilysine ER retention motif would be removed, and SEAP would be secreted. **d**) Implementation of AND logic by adding the N-terminal p450 signal anchor sequence as a second ER retention domain, so that both proteases would have to be present to remove both retention domains and allow SEAP to be secreted. **e)** A two-protease cascade was created where TEVP was required to activate TVMVP, which subsequently cleaved SEAP RELEASE. SEAP secretion increased when TEVP was expressed (right panel). Each dot represents an individual biological replicate. Mean values were calculated from three (**b**) or four replicates (**c-e**). Error bars represent +/- SEM. The results are representative of at least two independent experiments; significance was tested by one-way ANOVA with a Tukey’s post-hoc comparison test among the multiple conditions. * = p < 0.05, *** = p < 0.001, **** = p < 0.0001

In addition to the parallel regulation of multiple outputs, another useful capability is the integration of multiple inputs. Logic operation is crucial for integrating multiple signals, previously implemented for protease circuits using degrons^7^ or coiled-coiled (CC) dimerization domains^6^. RELEASE enables the compact implementation of Boolean logic directly at the retention level. To implement OR, two protease cut sites were inserted in tandem into the cytosolic linker so that the retention motif is removed by either protease (**Fig. 2c**). To implement AND, a RELEASE complex was created containing the N-terminal p450 signal anchor sequence and the C-terminal dilysine motif, each alone conferring sufficient ER retention (**Fig. 2d**). For SEAP to be secreted, both motifs must be removed (**Fig. 2d**). We attributed the reduced secretion in the AND gate construct due to the use of the N-terminal signal anchor sequence (**Supplementary Fig. 2b**), which confers retention by directly inserting into the ER membrane^36,37^ rather than retention through retrograde transport^31,32^. Both gates function as expected (**Fig. 2c, 2d**). We also implemented an alternative approach for AND (**Supplementary Fig. 4**)^45,46^.

Other than processing signals on its own, can RELEASE be coupled to other protease circuitsã We used protease-activated protease as an example of such circuits^6^. We used CC domains to associate split protease halves with complementary catalytically-inactive halves (**Fig. 2e**), “caging’ them by preventing the active halves from associating with each other. Cut sites were incorporated adjacent to (or within) the linker regions, allowing the input protease to remove the inhibitory domains. Following removal of the autoinhibitory portion, the complementary CC domains of the functional split protease halves would then associate and reconstitute protease activity (**Fig. 2e**). Using this approach, we created a two-protease cascade, in which TEVP activates TVMVP, which in turn cleaves the TVMVP-inducible RELEASE. This circuit increased SEAP secretion in response to TEVP, while maintaining strong retention in the absence of TEVP (**Fig. 2f**). This highlights the modularity of the RELEASE design and the ability to engineer additional functionality into it.

### RELEASE controls biologically relevant proteins

Many cytokines are pleiotropic and their systemic administration would cause serious adverse effects, so controlling their local expression with RELEASE would be advantageous for tumor immunotherapy^19^. We selected interleukin 12 p70 (referred to as IL-12), because it is a immunomodulatory cytokine important for T-cell activation and proliferation^47,48^. IL-12 is composed of two obligatory subunits (p35 and p40)^49^, so we fused the two subunits with a flexible linker^19,50^ and then with RELEASE (**Fig. 3a**). As expected, TVMVP significantly increases IL-12 secretion (**Fig. 3b**).

**Figure 3:**
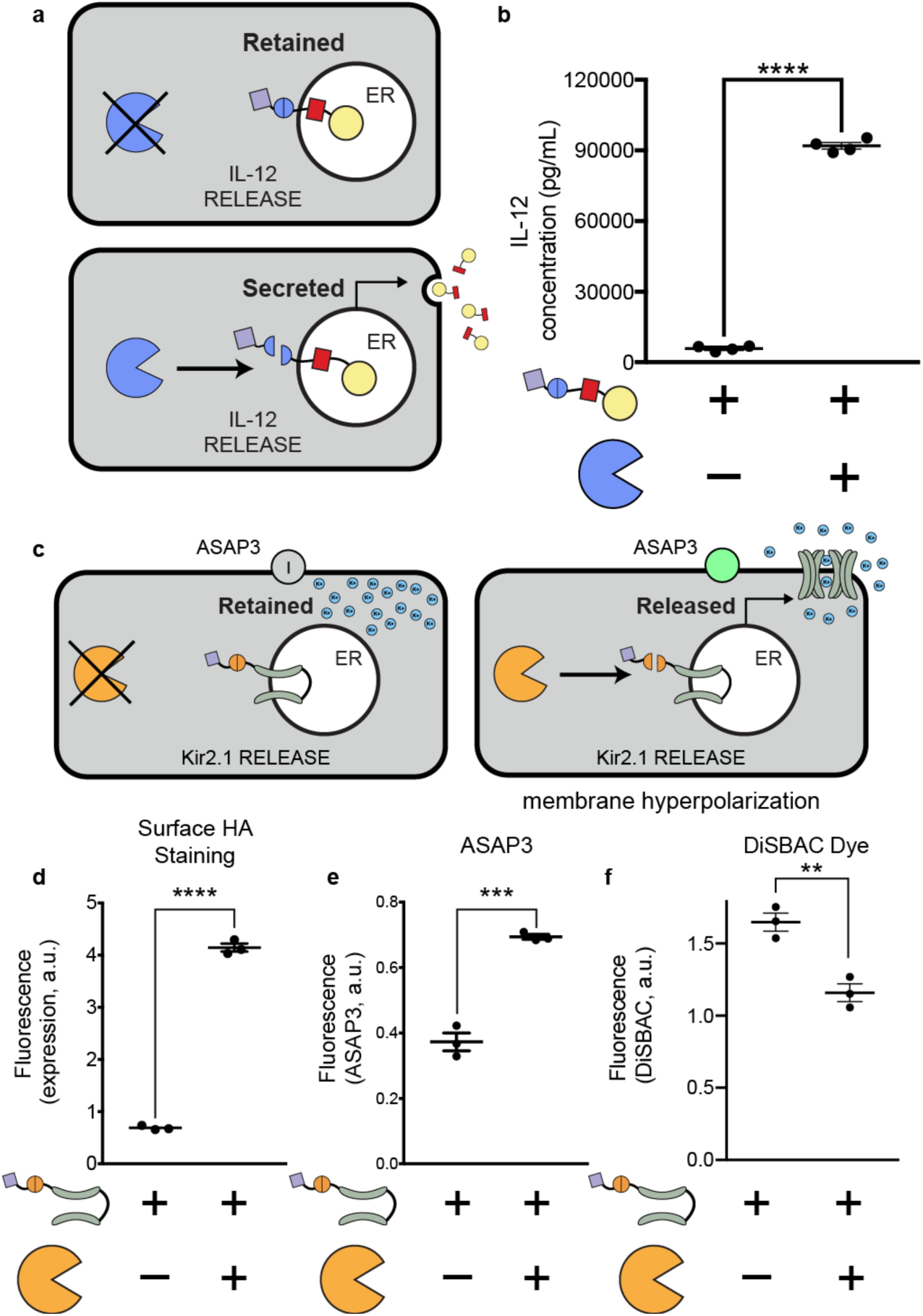
Controlling bioactive proteins using RELEASE. **a)** The cytokine IL-12 was fused to RELEASE and placed under the control of TVMVP. **b)** TVMVP significantly increase IL-12 secretion. **c)** The inwardly rectifying potassium channel Kir2.1 was fused to RELEASE. In addition, the genetically encoded voltage indicator ASAP3 was co-transfected. **d)** Co-expression of Kir2.1-RELEASE with TEVP resulted in a significant increase in the amount of Kir2.1 expressed on the surface, which was quantified using surface staining for HA and flow cytometry. The surface display of functional Kir2.1 in response to TVMVP was shown to cause hyperpolarization of transfected cells. This was validated by measuring change in the fluorescence intensity of the genetic reporter **e)** ASAP3, or the chemical dye, **f)** DiSBAC2(3). Each dot represents an individual biological replicate. Mean values were calculated from four (**b**) or three replicates (**d-f**). Error bars represent +/- SEM. The results are representative of at least two independent experiments. Significance was tested using an unpaired two-tailed Student’s *t*-test between the two indicated conditions for each experiment. ** = p < 0.01, *** = p < 0.001, **** = p < 0.0001

As for controlling membrane proteins, we chose the Kir2.1 potassium channel as an example of (**Fig. 3c**), because it is a powerful tool in neurobiology^51,52^ and a well-characterized model membrane protein. A protease-controlled Kir2.1 would enable the conditional silencing of neurons based on their intracellular states or extracellular cues, e.g., therapeutic silencing of the most active neurons during a seizure without the side effects of conventional methods that exert indiscriminate silencing. Unlike secreted proteins, Kir2.1 has cytosolic motifs that directs its transport in the secretory pathway, posing unique challenges for RELEASE and serving as a test case for its future adaptation to other membrane proteins. To measure the surface display of Kir2.1, a hemagglutinin (HA) epitope was incorporated into its extracellular loop^34^. Initial experiments fusing Kir2.1 with the standard RELEASE construct resulted in leaky display of Kir2.1 in the absence of TEVP (**Supplementary Fig. 5**). We reasoned that it is because Kir2.1 has a long cytosolic tail, and that the dilysine motif is the most effective when positioned closely to the ER membrane^30,34^. In contrast, another ER retention motif, RXR, is most effective when positioned distally from the membrane^34^. Indeed, a RELEASE construct using the RXR motif, improved retention (**Supplementary Fig. 5**), and successfully controlled its surface display using TEVP (**Fig. 3d**).

Kir2.1 functions as a homo-tetramer^53^, provoking the question of whether the RELEASE system could interfere with tetramerization and consequently channel function (**Fig. 3c**). Surface display of functional Kir2.1 leads to efflux of potassium ions and hyperpolarization^53^, providing a metric we can use to assess the its functionality. We used two reporters to measure changes in membrane potential: ASAP3^54^ and DiSBAC_2_(3)^55^. ASAP3 is a genetically encoded voltage indicator^54^ that increases fluorescence as cells become hyperpolarized, while DiSBAC_2_(3) is a chemical dye that decrease cell entry and therefore fluorescence intensity upon hyperpolarization^56^. When Kir2.1 RELEASE was co-expressed with ASAP3, we observed a significant increase in fluorescence intensity in response to TEVP (**Fig. 3e**), suggesting Kir2.1 was functional. The chemical dye DiSBAC_2_(3) showed similar results (**Fig. 3f**), and the observed change in median fluorescent intensity was indicative of a 30 mV change in membrane potential^55^. Thus RELEASE-regulated Kir2.1 maintains its functionality.

### RELEASE responds to oncogenic inputs

One of the most compelling cases for protein circuits is therapy against recalcitrant cancers. The RAS family of proteins (HRAS, KRAS, and NRAS) provide a remarkable example.^57,58^. The activating RAS mutations have been implicated in a multitude of hard-to-treat cancers such as pancreatic ductal adenocarcinoma^57–59^ and non-small lung cancer^60^. The pharmacological targeting of RAS has been challenging^61–63^. We envision a “circuit as medicine” alternative, where an intracellularly introduced circuit interrogates the cancerous state of a cell, and conditionally lyses RAS-mutant cells, while programming cytokine secretion to activate a broader local immune response.

As a first step towards that vision, we hypothesized that we could exploit protein interaction during RAS signaling to activate RELEASE. RAS resides in the cell membrane^64,65^, and activated RAS recruits to the membrane effector proteins such as Raf^65–67^. To sense active RAS, we fused the N- and C-terminal halves of split TEVP to the RAS-binding domain (RBD) of Raf (**Fig. 4a**). The increased local concentration of the RBD-split TEVP sensor in response to activated RAS, along with their transition from the 3D cytosol to the more restrictive 2D membrane, was expected to facilitate the association of the protease halves through their residual mutual affinity^64^.

**Figure 4:**
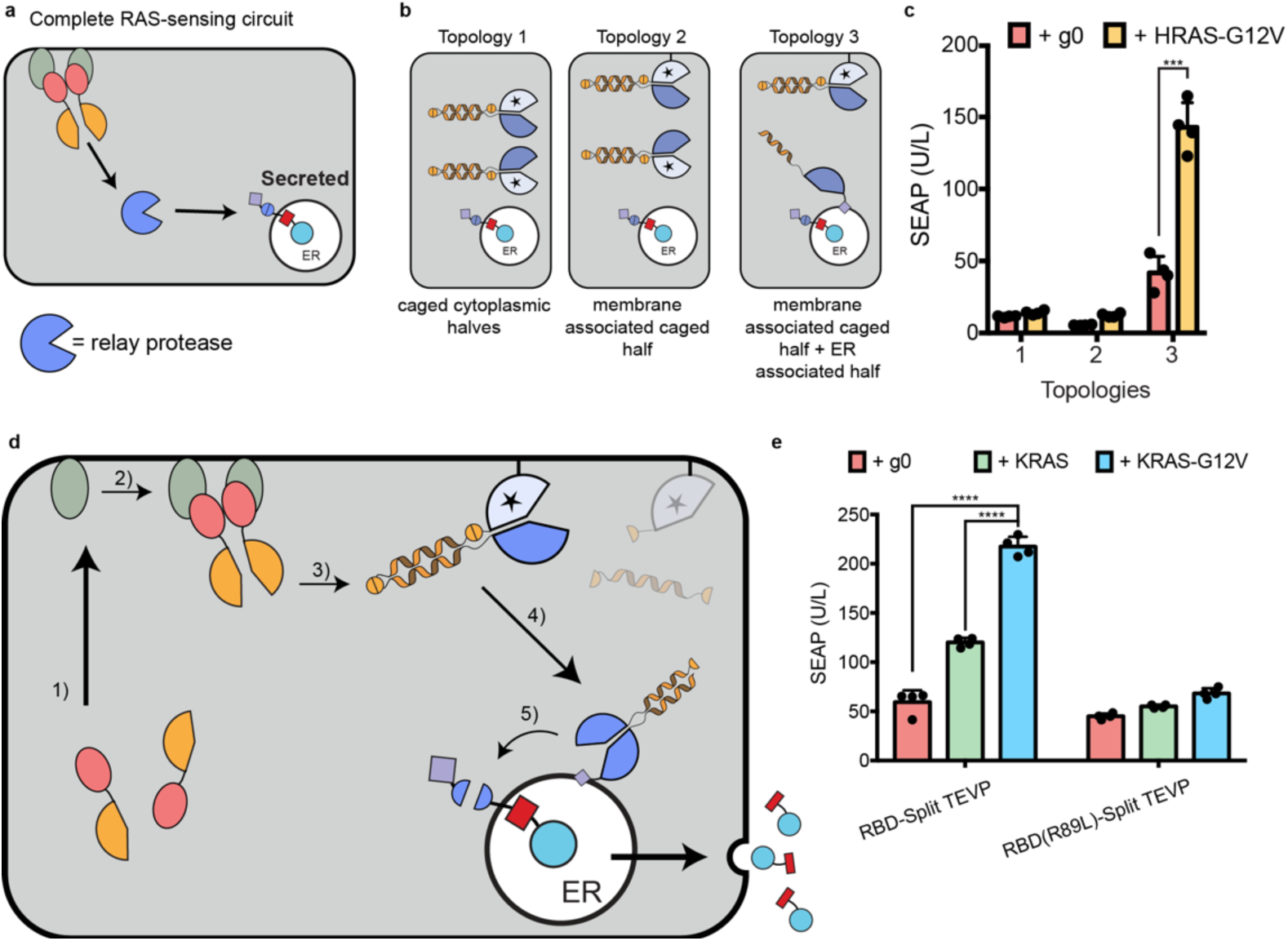
RAS-sensing circuit and protease replaying pathways to activate RELEASE. **a)** To sense active RAS, split TEVP was fused to the RBD domain of c-RAF. RBD-split TEVP binds to active RAS at the membrane surface of the cell where the two protease halves reassociated and reconstituted protease activity. Protease activation is propagated through an intermediate protease to relay the information from the cell membrane to the ER. **b)** Using protein localization motifs, three different topologies of intermediate protease components were created. Topology 1 uses two caged intermediate TVMVP protease halves in the cytosol. Topology 2 uses the same caged intermediate TVMVP, but with one half of the active protease localized to the membrane. Finally, Topology 3 has one half of the intermediate protease associated with the membrane, and the other half uncaged and present at the ER membrane via the p450 signal anchor sequence. The CC domain present on the uncaged TVMVP half (that was associated with the membrane) drives association with the complementary TVMVP half at the ER. **c)** There was a significant difference in the amount of SEAP secreted when using intermediate protease topology 3, with and without mutant HRAS-G12V, compared to topologies 1, and 2. **d)** Schematic of the signal processing of the complete KRAS-sensing circuit. The complete RAS-sensing circuit was activated by RBD-split TEVP interacting with active KRAS-G12V (1). The reconstituted TEV (2) then uncaged the membrane associated split TVMVP, releasing it from the membrane (3). The uncaged TVMVP contains a CC domain, which drives its association with the complementary CC domain present on the other split TVMVP half anchored to ER membrane (4). Finally, the reconstituted TVMVP cleaves the ER retention motif of RELEASE to secrete SEAP (5**). e)** Using the complete RAS-sensing circuit, we observed a significant increase in SEAP secretion when expressing an active mutant variant KRAS-G12V relative to baseline levels, or wildtype KRAS. This difference was not observed when using an RBD-Split TEVP containing the R89L mutation that reduced the association with active KRAS. Each dot represents an individual biological replicate. Mean values were calculated from four replicates (**c, e**). The error bars represent +/- SEM. The results are and representative of at least two independent experiments. Significance was tested using an unpaired two-tailed Student’s *t*-test between the two indicated conditions for each experiment. ** = p < 0.01, *** = p < 0.001, **** = p < 0.0001.

Building on our previous constructs sensing the RAS pathway^7^, we performed initial experiments using HRAS-G12V and the RBD-split TEVP sensor, and observed a minimal increase of SEAP secretion when regulated by TEVP-responsive RELEASE (**Supplementary Fig. 6**). Since HRAS-G12V reconstitutes RBD-split TEVP at the cell membrane, and cleavage of RELEASE occurs at the ER, we hypothesized that additional protease components would be required to propagate the signal from the cell membrane to the ER (**Fig. 4a**). Using the caged TVMVP intermediate protease (**Fig. 2d, 4b – topology** 1) did not improve SEAP secretion in response to HRAS-G12V, so we further hypothesized that spatial localization of the intermediate protease might be required to increase signal transduction. We first tried to increase the cleavage of the intermediate protease by bringing it closer to the TEVP input, fusing the C-terminal membrane transfer CAAX motif^68^ (**Supplementary Fig. 7a – left panel)** to one half of the caged split TVMVP (**Fig. 4b – topology 2**), but this did not improve SEAP secretion (**Fig. 4c**). We then also increased the possibility for the reconstituted intermediate protease to activate RELEASE, by fusing the uncaged other half of TVMVP with the signal anchor sequence of cytochrome p450 and therefore targeting it to the ER membrane (**Fig. 4b – topology 3**). This resulted in the greatest SEAP secretion in response to HRAS-G12V (**Fig. 4c**). After titrating down the ER-bound uncaged half of TVMVP, we reduced background and improved dynamic range (**Supplementary Fig. 7b**).

We then generalized the design to KRAS, the most frequently mutated RAS in cancer^69^. We validated that our circuit responds very similarly to KRAS-G12V and HRAS-G12V (**Supplementary Fig. 7c**), probably because RAS isoforms share up to 90% homology in the region where RBD binds^61,70^. As a control, the split TEVP sensor fused to the RBD mutant (R89L), which has a reduced affinity to activated RAS^64,67^, did not significantly increase SEAP secretion in response to HRAS-G12V or KRAS-G12V (**Fig. 4e**).

We reasoned that the choice of cell membrane-localization domains might affect baseline, because post-translational modification of CAAX initially inserts the protein at the ER membrane^71^, which could facilitate TVMVP reconstitution in the absence of TEVP inputs. To further reduce the background of the RAS sensor, we additionally tested the N-terminal membrane anchoring portion of the SH4 domain of Lyn and Fyn tyrosine kinases^72^, the cell membrane-targeting of which bypasses ER^72^. The Lyn and Fyn motifs reduced background SEAP secretion relative to the CAAX motif (**Supplementary Fig. 7d**), and enabled increased SEAP secretion without significantly increasing the background (**Supplementary Fig. 7e**).

The complete circuit is summarized in **Figure 4d**. We observed that the circuit was responsive to the oncogenic state of KRAS, since cells secreted significantly more SEAP when co-expressed with active mutants of KRAS (**Fig. 4e – blue bar, Supplementary Fig. 7g**) compared to wildtype KRAS (**Fig. 4e – green bar**), and endogenous wildtype KRAS (**Fig. 4e – red bar**). The oncogenic state of KRAS also resulted in a much smaller and statistically insignificant increase in SEAP secretion when using the RBD-split TEVP R89L mutant (**Fig. 4e**).

### Plug-and-play capabilities of RELEASE

In addition to building towards RAS detection, our RAS-centric engineering efforts also established a plug-and-play protein circuit framework. RELEASE, in conjunction with CHOMP and other protease components, enables the detection of any input that can be converted to dimerization or proteolysis. This signal can then be processed by RELEASE itself or other protease circuits to control the display or secretion of proteins (**Fig. 5a**).

**Figure 5:**
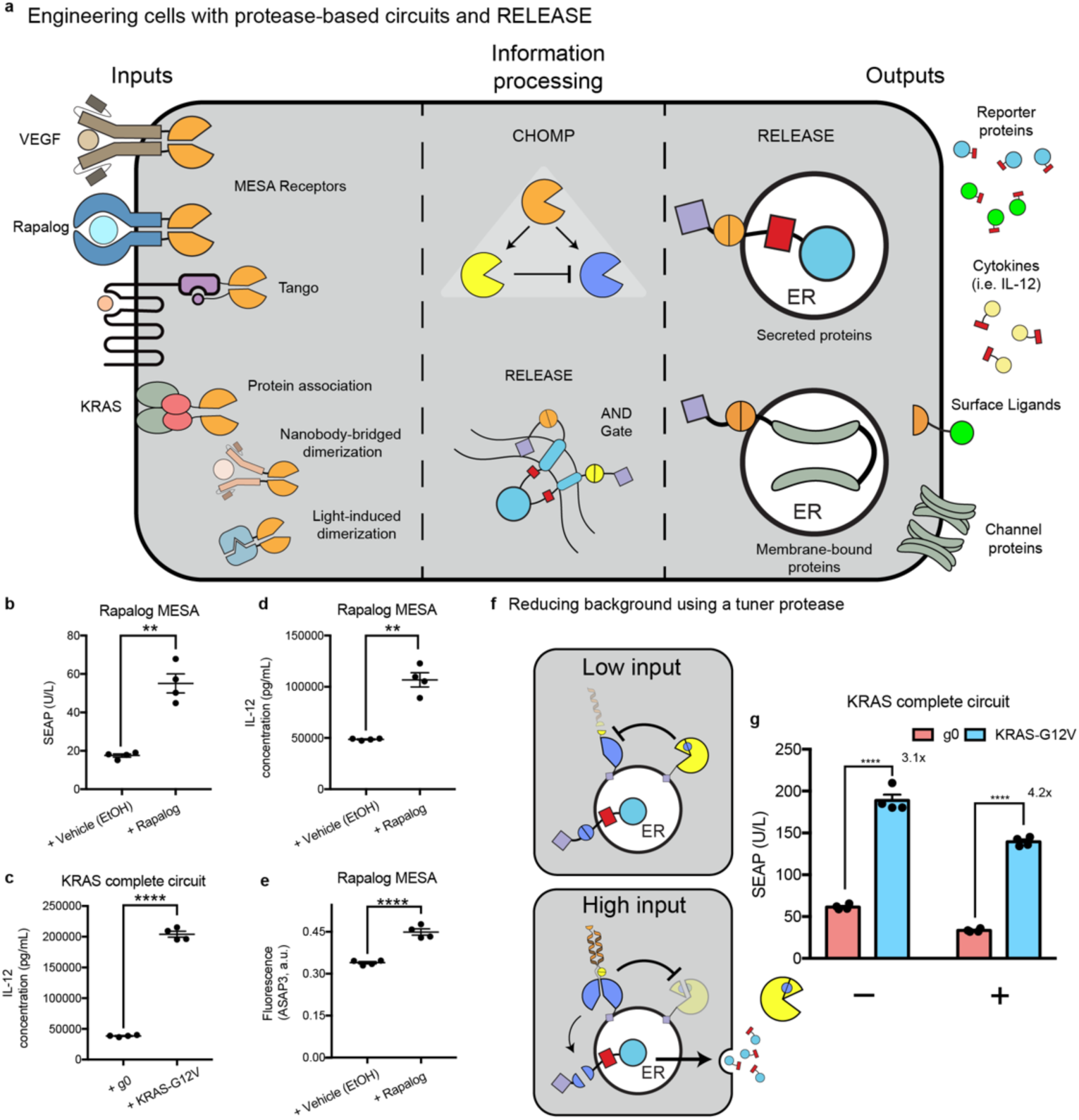
Plug-and-play capabilities of RELEASE. **a)** Any multimerization event, such as ligand-induced receptor dimerization (i.e. MESA receptors, or Tango), protein association, nanobody-bridged dimerization, or light-induced dimerization can be harnessed to reconstitute and activate split proteases. This information can then be processed using CHOMP circuits or even RELEASE itself to produce complex responses. Each component of the engineered can be optimized independently of each other and are not necessarily dependent on the input or output components. To highlight the plug-and-play capabilities of RELEASE, we tested different input and output combinations, while keeping the intermediate CHOMP circuit intact. **b)** Using the rapalog MESA receptor as the input, SEAP secretion was controlled. IL-12 secretion was induced by **c)** KRAS expression or induction with **d)** rapalog. **e)** We also observed Kir2.1-mediated hyperpolarization after induction with Rapalog. **f)** Schematic of CHOMP circuit containing reciprocal inhibition of TVMVP and HCVP to reduce background activity of RELEASE. When the amount of input is low, the ER-associated split TVMVP protease is repressed by the ER-associated HCVP through removing the complementary CC motif, reducing the association with the other split functional half. When the amount of input is high, fully reconstituted TVMVP will be present at higher levels and repress HCVP by removing the core HCVP from its activity-enhancing co-peptide (small yellow pie space). **g)** Addition of the tuner protease increased the dynamic range of the RAS-sensing circuit, by reducing baseline secretion. Each dot represents an individual biological replicate. Mean values were calculated from four biological replicates (**b-e, g**). Error bars represent +/- SEM. The results are representative of at least two independent experiments. Significance was tested using an unpaired two-tailed Student’s *t*-test between the two indicated conditions for each experiment. ** = p < 0.01, **** = p < 0.0001.

As a proof of principle, we used the well-established MESA receptor (membrane-localized split TEVP reconstituted by rapalog)^25,26,73^ as an input to activate RELEASE via the intermediate protease circuit optimized above (**Fig. 4c**). Switching the input components to the rapalog MESA receptor, we increased SEAP secretion in response to rapalog (**Fig. 5b**). We also used RELEASE to control the secretion of IL-12 in response to mutant KRAS (**Fig. 5c**) or rapalog (**Fig. 5d**), and to control the surface display of Kir2.1 by rapalog (**Fig. 5e**).

The processing protease circuit is also modular. Specific applications of RELEASE may require a greater dynamic range or more complex dynamic secretion patterns that can be achieved by incorporating additional orthogonal proteases^6,7^. For example, to improve the dynamic range of the RAS-sensing circuit, we incorporated a previously established positive feedback loop based on reciprocal inhibition between TVMVP and HCVP to tune the level TVMVP^7^ (**Fig. 5f**). When input was low, or not present, HCVP would inactive the “baseline” reconstitution of TVMVP by removing the complementary CC domain (**Fig. 5f – top panel**). However, when there was sufficient input (KRAS-G12V^+^ cells), the reconstituted TVMVP would override HCVP by removing its activating co-peptide (**Fig. 5f – bottom panel**). By varying the amount of HCVP transfected, we reduced the background activity and increased the dynamic range of the engineered cells containing the complete RAS circuit (**Fig. 5g**). These results demonstrate the possibly of tuning RELEASE with additional proteases and eventually creating more complex responses.

## Discussion

Here, we introduced the generalized protease-responsive platform RELEASE to control the secretion and display of proteins (**Fig. 1**). RELEASE is compatible with protein-level circuit operations (**Fig. 2)**, and enables plug-and-play control of various outputs (Figs. 3, 5) using a variety of inputs (Fig. 4, 5). For all these examples, we simply switched the input and output (RELEASE) components, while keeping the intermediate protease chassis intact – without any re-optimization. This highlights the modularity of using protease-based sensors, protease circuits, and RELEASE to engineer sense-and-response capabilities.

When adapting RELEASE for new applications, all one needs is a protein-mediated dimerization event that could be harnessed to reconstitute protease activity (**Fig. 4a, 5a**). We can therefore tap into additional synthetic receptors platforms that rely on ligand-induced dimerization, such the Generalized Extracellular Molecule Sensor (GEMS)^42^, or Tango^74^. In this work we demonstrate that one can use intermediate proteases to propagate protease signal from the cell membrane to the ER to activate RELEASE (Fig. 4a), suggesting that using alternative motifs may allow for signal propagation from other subcellular locations, such as nucleus or mitochondria, to ER. Because the components of the conventional protein secretion pathway are conserved among different cell types and species, we expect RELEASE to function in these different contexts as well.

RELEASE enables novel therapeutic modalities. For example, we hypothesize that we can use the KRAS-sensing circuit (**Fig. 4c**) to selectively express immunostimulatory signals (such as IL-12, surface T-cell engagers, and anti-PD1) to mark cancer cells for T-cell mediated destruction without affecting normal cells^19^. The selectivity of the circuit will be further improved using additional proteases through quantitative thresholding (**Fig. 5g**) or logic operations. For the latter, many RAS-driven cancers harbour additional mutations to tumor suppressor proteins, such as p53^75^. One could use split proteases fused to nanobodies^4,7^ that have preferential binding to mutant p53^75^, to activate RELEASE only when both mutant KRAS and mutant p53 are simultaneously present, via AND logic (**Fig. 2c**). An additional benefit is that the protease circuit components can be encoded within single mRNA transcripts^76^ that do not pose the risk of insertional mutagenesis.

RELEASE will also expedite other potential therapeutic applications in fields as diverse as neurobiology, developmental biology, immunology, tissue engineering, and transplantation, to name a few. To take a third and last example, in addition to the cancer immunotherapy and neuronal silencing applications discussed above, RELEASE can be used to create sense-and-respond cells to control immunomodulating cytokines and growth factors important for graft acceptance, such as IL-10^77^ and TGF-β^20^, which cannot normally be delivered systemically due to their pleiotropic and off-target effects. Co-delivering these engineered cells with therapeutic cells, such as pancreatic islets may be a suitable approach create engineered tissue implants that can engraft without the need for systemic immunosuppression. The proposed plug-and-play sense and secretion components using RELEASE would allow for the programming of such communications with unprecedented specificity and precision.

## Acknowledgments

We would like to thank Dr. Lin, and Dr. Leonard for kindly sharing some of their plasmids that were used in this work. We would also like to thank Leo Scheller for providing the protocol for the SEAP assays. This work was funded by NIH (4R00EB027723-02, X.J.G), Stanford Cancer Institute (Cancer Innovation Award, X.J.G.), Stanford SystemX Alliance (Seed Grant, X.J.G.), NSERC (PDF-557516-2021, A.E.V.), the International Human Frontier Science Program Organization (LT000221/2021-L, A.E.V.), and the Stanford Graduate Fellowship (J. K. and C. A.).

## Author Contributions

A.E.V. and X.J.G. conceived and directed the study. A.E.V, J.K, and C.A. performed most of the experiments. X.J.G. created the HRAS-sensing protease, and L.S.C and R.Z. created the protease-activated protease under M.B.E.’s supervision. A.E.V, and J.K. analysed the data for the manuscript. A.E.V., J.K. and X.J.G. wrote the manuscript. All authors provided feedback on the manuscript.

## Materials and Methods

### Plasmid generation

All plasmids were constructed using general practices. Backbones were linearized via restriction digestion, and inserts were generated using PCR, or purchased from Twist Biosciences. MESA-rapalog receptor source plasmids were a generous gift from Joshua Leonard^26^. The plasmid containing the voltage indicator, ASAP3 was a generous gift from Michael Lin^54^. A complete list of plasmids used in this study can be found in **supplementary table 2**, and all maps will be deposited on Addgene after publication.

### Tissue culture

Flp-In™ T-REx™ Human Embryonic Kidney (HEK) 293 cells were purchased from Thermo Scientific (Catlog# R78007). Cells were cultured in a humidity-controlled incubator under standard culture conditions (37°C with 5% CO_2_) in Dulbecco’s Modified Eagle Medium (DMEM), supplemented with 10% fetal bovine serum (FBS - Fisher Scientific; catalog# FB12999102), 1 mM sodium pyruvate (EMD Millipore; catalog# TMS-005-C), 1X Pen-Strep (Genesee; catalog# 25-512), 2 mM L-glutamine (Genesee, catalog# 25-509) and 1X MEM non-essential amino acids (Genesee; catalog# 25-536). To induce expression of transiently transfected plasmids, 100 ng/mL of Doxycycline was added at the time of transfection. Rapalog AP21967 (also known as A/C heterodimerizer, purchased from Takara Biosciences; catalog# 635056) is a synthetic rapamycin analog that can bind with FRB harboring the T2098L mutation, and is designed not to interfere with the native mTOR pathway^78^. All our constructs in this study using the FRB protein contain the T2098L mutation and were induced with 100 nM of rapalog, unless otherwise stated.

### Transient transfections

HEK 293T cells were cultured in either 24-well or 96-well tissue culture-treated plates under standard culture conditions. When cells were 70-90% confluent, the cells were transiently transfected with plasmid constructs using the jetOPTIMUS® DNA transfection Reagent (Polyplus transfection, catalog# 117-15), as per manufacturer’s instructions.

### Measuring protein secretion

Secreted Alkaline Phosphatase (SEAP) Assay was performed as previously described^42^. Briefly, following two days after transient transfection, the supernatant was collected without disrupting the cells and heat inactivated at 70°C for 45 minutes. Following heat inactivation, 10-40 μL of the supernatant was mixed with dH_2_O for a final volume of 80 μL, and then mixed with 100 μL of 2X SEAP buffer (20 mM homoarginine (ThermoFisher catalog# H27387), 1 mM MgCl_2_, and 21% (v/v) dioethanolamine (ThermoFisher, catalog# A13389)) and 20 uL of the p-nitrophenyl phosphate (PNPP, Acros Organics catalog# MFCD00066288) substrate (120 mM). Samples were measured via kinetic measurements (1 measurement/min) for a total of 30 minutes at 405 nm using a SpectraMax iD3 spectrophotometer (Molecular Devices).

Secreted GFP was measured by incubating cell-free supernatant with cells displaying the Gbp6 GFP-binding nanobody, with mCherry fused to its cytosolic tail as a co-transfection marker. Captured GFP was used to quantify changes in the amount of secreted GFP in response to protease expression.

To measure the amount of secreted IL-12, cell-free supernatant was collected and quantified using the Human IL-12p70 DuoSet ELISA (R&D Systems; catalog# DY1270), as per the manufacturer’s instructions.

### Flow Cytometry and data analysis

Two days after transient transfection, cells were harvested using FACS buffer (HBSS + 2.5 mg/mL of Bovine Serum Albumin (BSA)). For experiments requiring antibody staining, surface GFP was measured by incubating cells with a 1:1000 dilution of anti-GFP Dylight 405 antibody (ThermoFischer; catalog# 600-146-215) in FACS buffer for one hour at 4°C. For experiments measuring the surface display of Kir2.1, cells were incubated with 1:500 dilution of anti-hemagglutinin antibody (HA, Abcam; catalog# ab137838), followed by incubation with a donkey anti-rabbit IgG conjugated to alexa-647 (Abcam, Cat# ab150075). After staining, cells were washed twice with FACS buffer and then strained using a 40 μm cell strainer. Cells were analyzed by flow cytometry (BioRad ZE5 Cell Analyzer). As previously described^7^, we use the EasyFlow Matlab-based software package developed by Yaron Antebi to process the flow cytometry data.

For analysis, we selected and compared cells with the highest expression of the co-transfection marker, which was typically mCherry. This was done to have the largest separation between basal reporter autofluorescence from cellular autofluorescence, as previously described^7,26^. For experiments using the Kir2.1 potassium channel, cells were either co-transfected with the voltage indicator ASAP3^54^ or incubated with the Oxonol chemical dye, DiSBAC_2_(3) as previously described^55^. The N-terminus of Kir2.1 was fused with mCherry, which acted as a co-transfection marker. After gating on cells with high expression of Kir2.1, the median fluorescence intensity was used to estimate changes in membrane potential^55^.

### Statistical analysis

Values are reported as the means from at least 3 biological replicates, which was representative from two independent biological experiments. For experiments comparing two groups, an unpaired Student’s *t*-test was used to assess significance, following confirmation that equal variance could be assumed (*F*-test). If equal variance could not be assumed, then a Welch’s correction was used. For experiments comparing three or more groups, a one-way ANOVA with a post hoc Tukey test was used to compare the means among the different experimental groups. Data were considered statistically significant at a p value of 0.05. Data are presented as average ± SEM, unless otherwise stated. All statistical analysis was performed using Prism 7.0 (GraphPad).

## Data Availability

All data reported in this paper are available from the corresponding author on request.

## Supplementary Files

### Supplementary Figure Captions

**Supplementary Figure 1:**
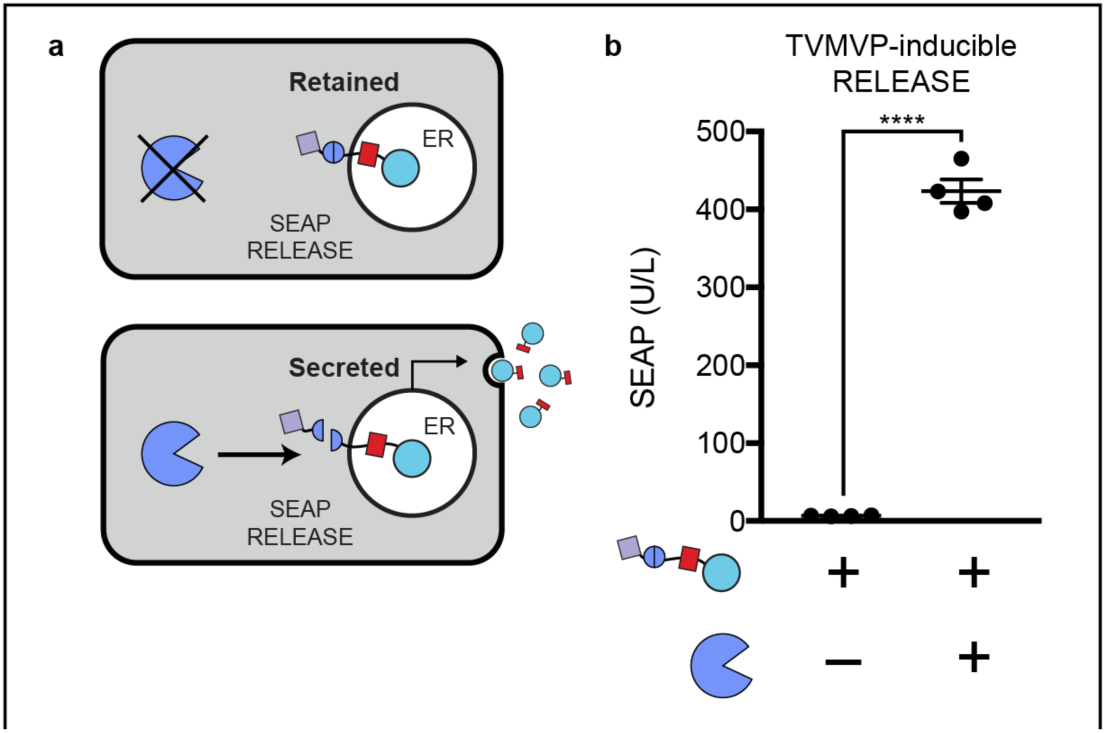
TVMVP-inducible RELEASE. **a)** Schematic of TVMVP-inducible RELEASE for controlling protein secretion. **b)** SEAP was fused to a TVMVP-inducible RELEASE, and co-expression with TVMVP secreted more SEAP. Each dot represents a biological replicate. Mean values were calculated from four biological replicates (**b**) +/- SEM. The results are representative of at least two independent experiments; significance was tested using an unpaired two-tailed Student’s *t*-test between the two indicated conditions. **** = p < 0.0001.

**Supplementary Figure 2:**
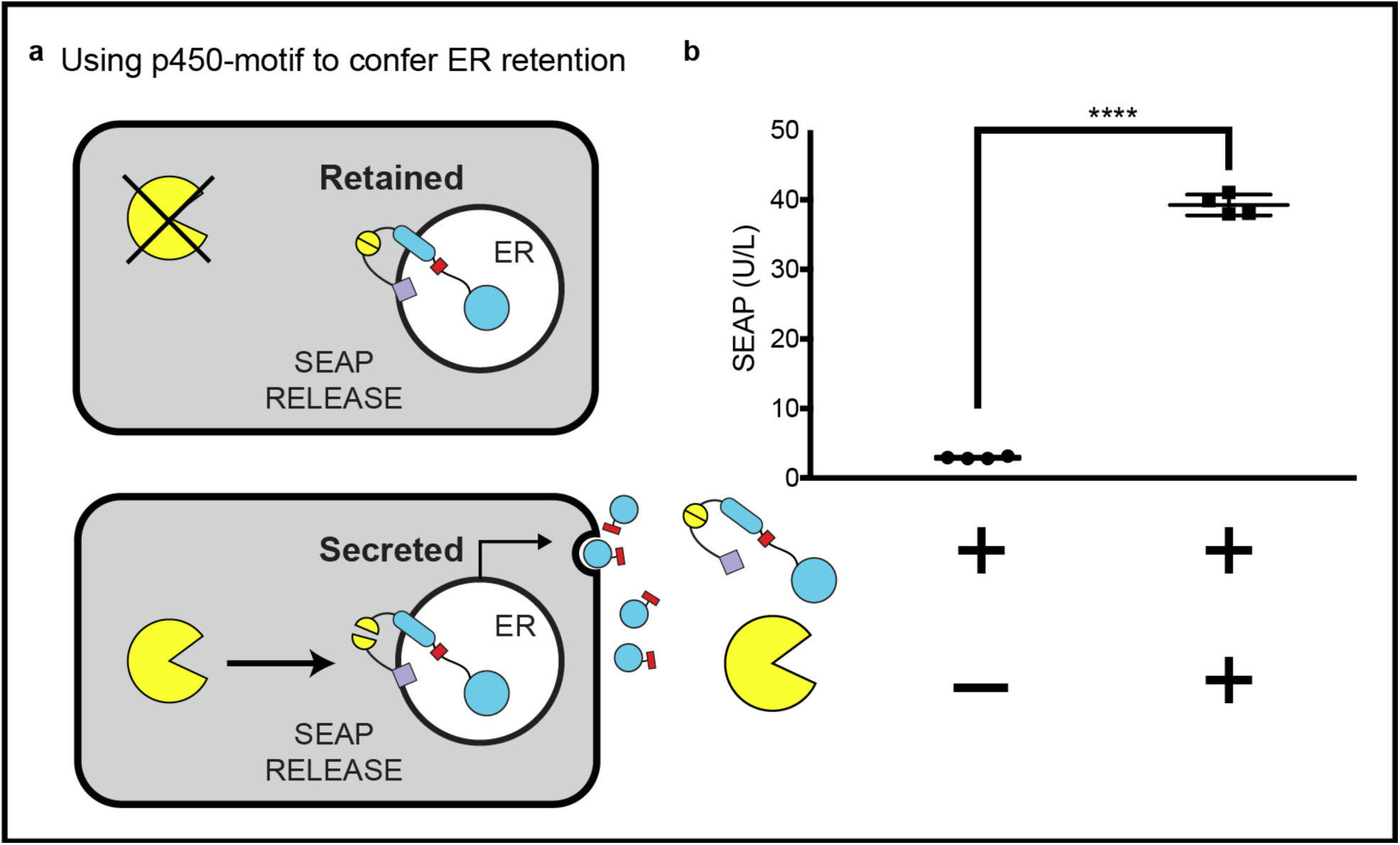
An alternative ER-retention domain to create RELEASE. **a)** Schematic of RELEASE using the N-terminal signal anchor sequence of cytochrome p450 to control protein secretion. **b)** When co-expressed with HCVP, SEAP secretion increased relative to when the HCVP was absent. Each dot represents a biological replicate. Mean values were calculated from four biological replicates (**b**) +/- SEM. The results are representative of at least two independent experiments; significance was tested using an unpaired two-tailed Student’s *t*-test between the two indicated conditions for each experiment. **** = p < 0.0001.

**Supplementary Figure 3:**
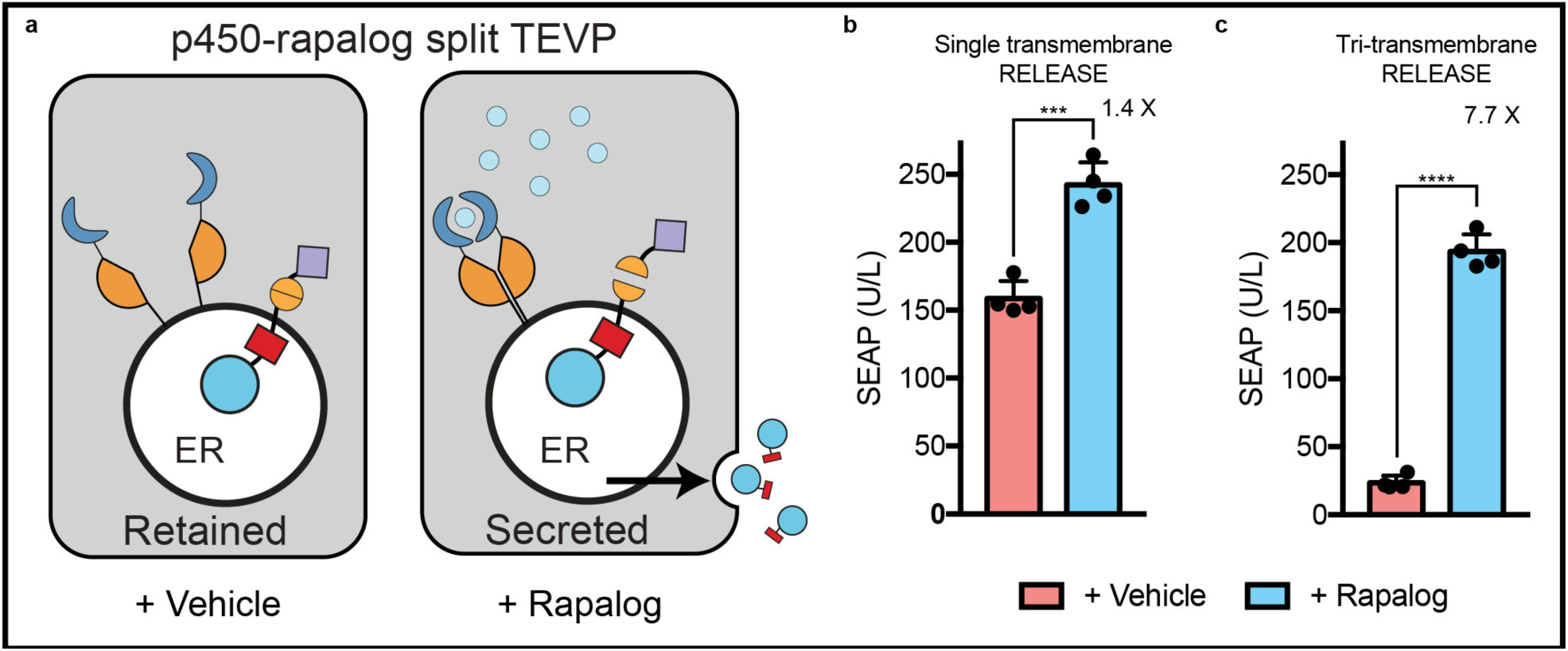
Dynamic range of RELEASE is increased by using different RELEASE constructs that have different cleavage efficiencies. **a)** Schematic of rapalog-inducible split TEVP localized to the ER membrane via the p450 signal anchor sequence. When rapalog is present, split TEVP will be reconstituted to cleave SEAP RELEASE. **b)** With the single transmembrane RELEASE construct, there was a minor increase in the SEAP secretion after induction with rapalog relative to the control. **c)** Using the tri-transmembrane RELEASE construct there was a greater difference in SEAP secretion compared to the single transmembrane RELEASE construct (7.7-fold vs. 1.4-fold). The difference between the fold-changes was attributed to the reduction in SEAP secretion under basal conditions with the tri-transmembrane RELEASE. Each dot represents a biological replicate. Mean values were calculated from four biological replicates (**b, c**) +/- SEM. The results are representative of at least two independent experiments; significance was tested using an unpaired two-tailed Student’s *t*-test between the two indicated conditions for each experiment. *** = p < 0.001, **** = p < 0.0001

**Supplementary Figure 4:**
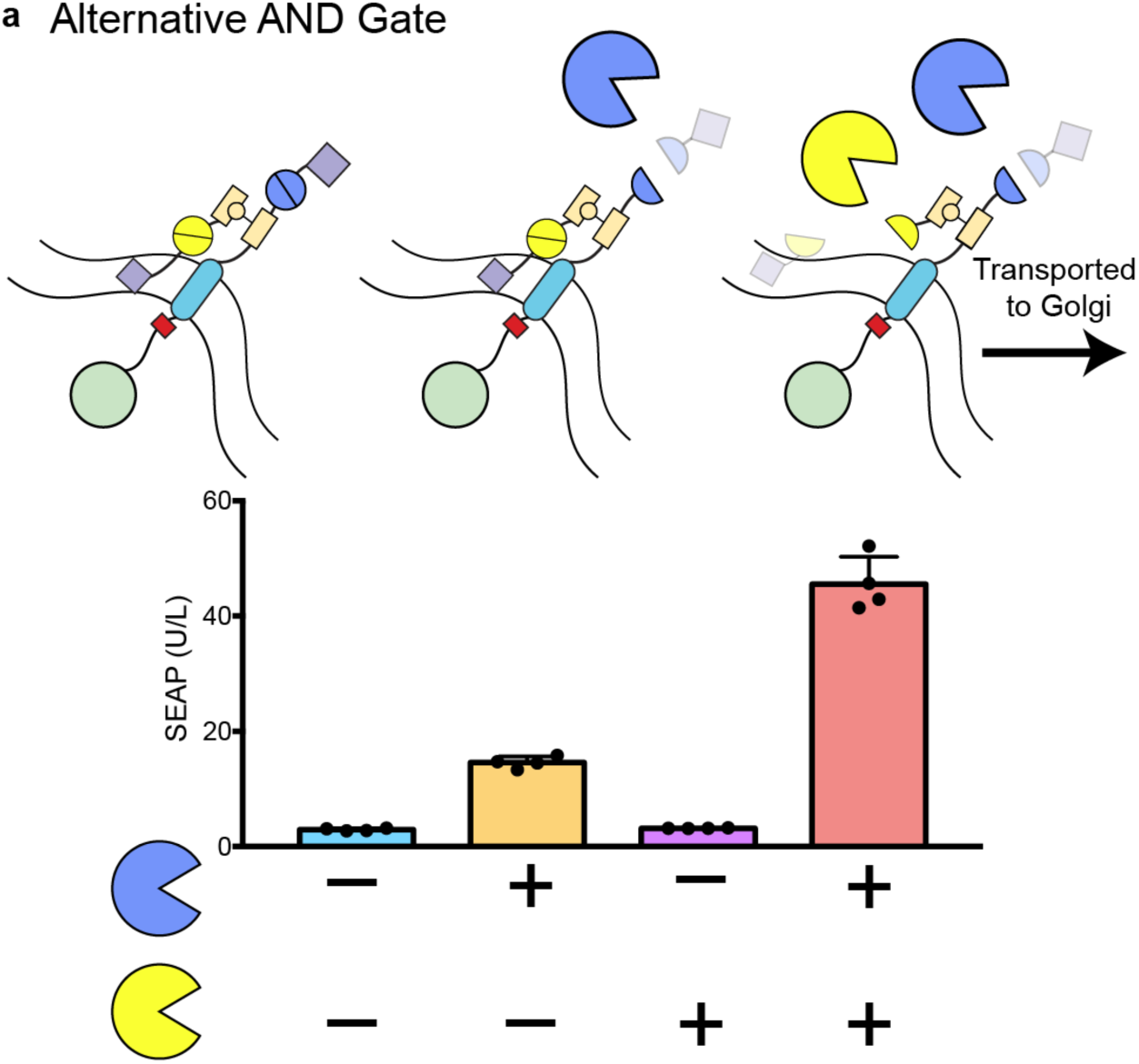
An alternative AND gate was implemented using the SpyTag/SpyCatcher peptide-protein pair. HCVP inducible SpyCatcher was localized to the ER using the signal anchor sequence of p450. A TVMVP inducible RELEASE construct containing an internal SpyTag peptide within the cytoplasmic linker region rapidly associated with the ER-retained SpyCatcher. SEAP secretion was dependent on the expression of both HCVP and TVMVP, however some SEAP was secreted when co-expressing TVMVP alone, which may be due to an incomplete reaction with SpyCatcher. Each dot represents a biological replicate. Mean values were calculated from four biological replicates (**b**) +/- SEM. The results are representative of at least two independent experiments.

**Supplementary Figure 5:**
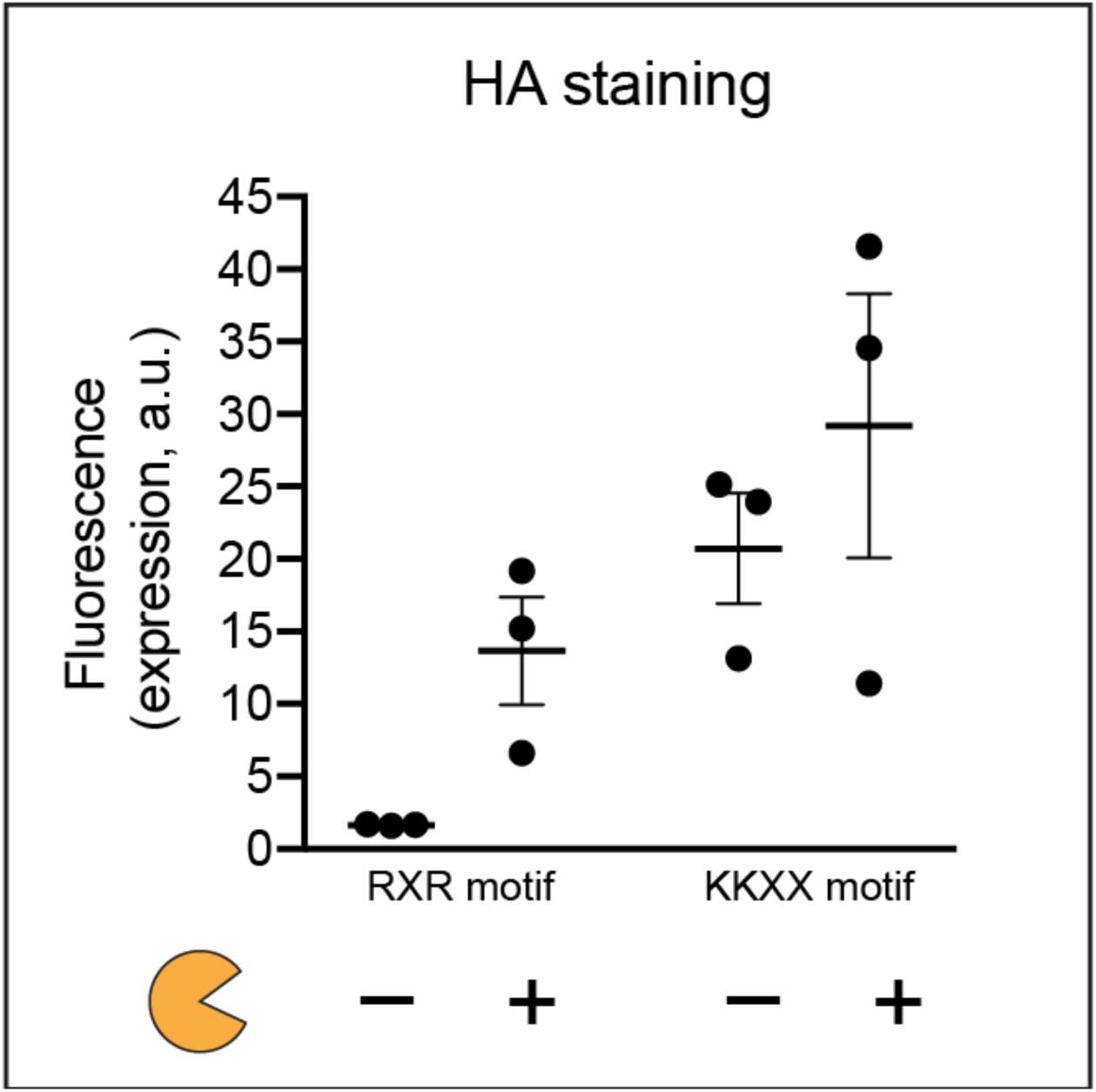
Surface display of Kir2.1 was dependent on the ER retention motif used in the RELEASE construct. Due to the large cytoplasmic tail of Kir2.1, the C-terminal was farther away from the ER membrane relative to other RELEASE constructs. The RXR motif retains proteins better than the KKXX motif when the C-terminal is distal to ER membrane. Each dot represents a biological replicate. Mean values were calculated from four biological replicates (**b**) +/- SEM. The results are representative of at least two independent experiments.

**Supplementary Figure 6:**
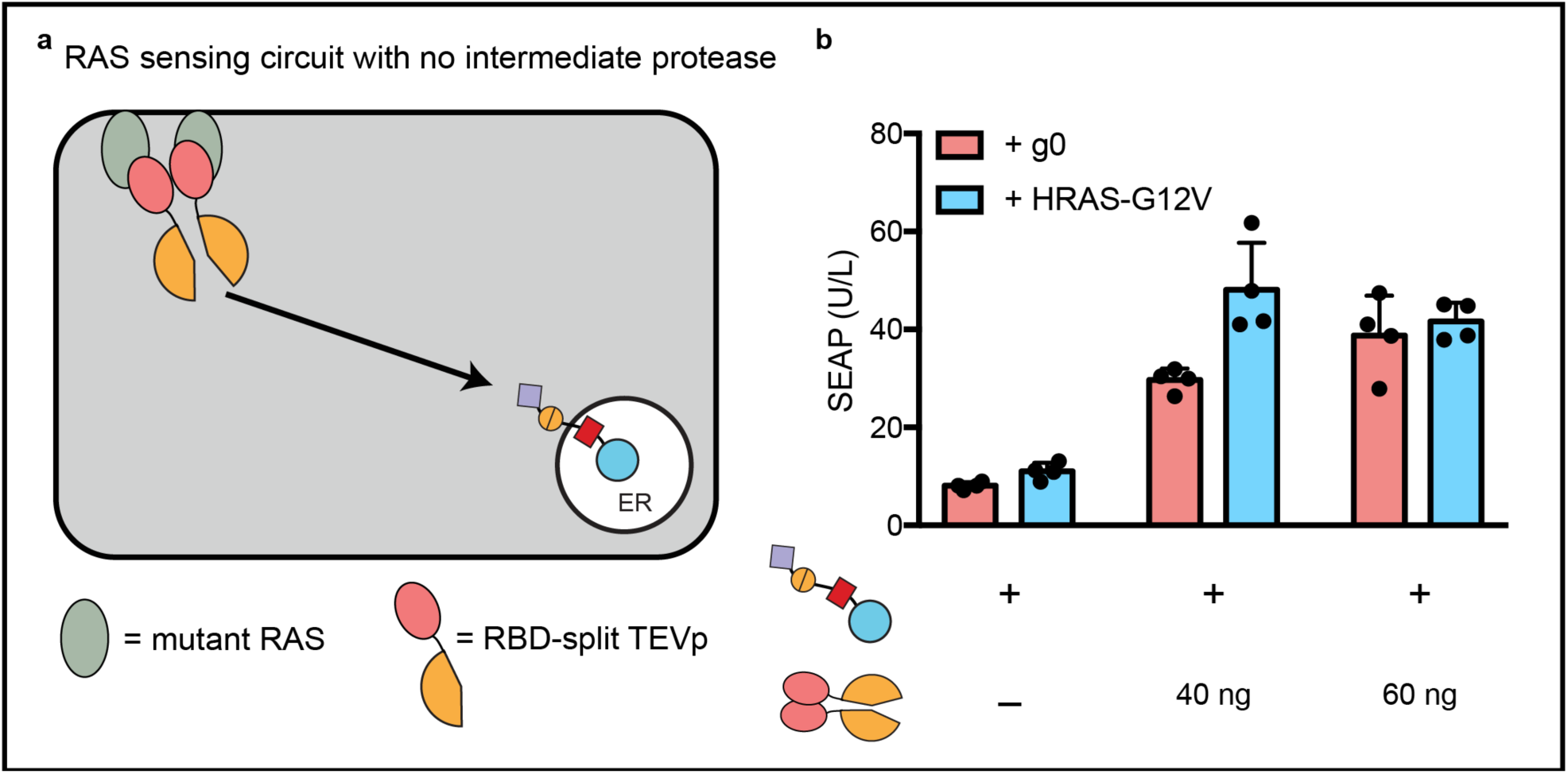
**a)** Schematic of RAS-sensing circuit without using intermediate protease to propagate the signal. **b)** The sensing of active mutant HRAS-G12V at the membrane using RBD-split TEVP did not result in a significant increase in the amount of SEAP secretion, relative to cells not containing mutant HRAS. Each dot represents a biological replicate. Mean values were calculated from four biological replicates (**b**) +/- SEM. The results are representative of at least two independent experiments.

**Supplementary Figure 7:**
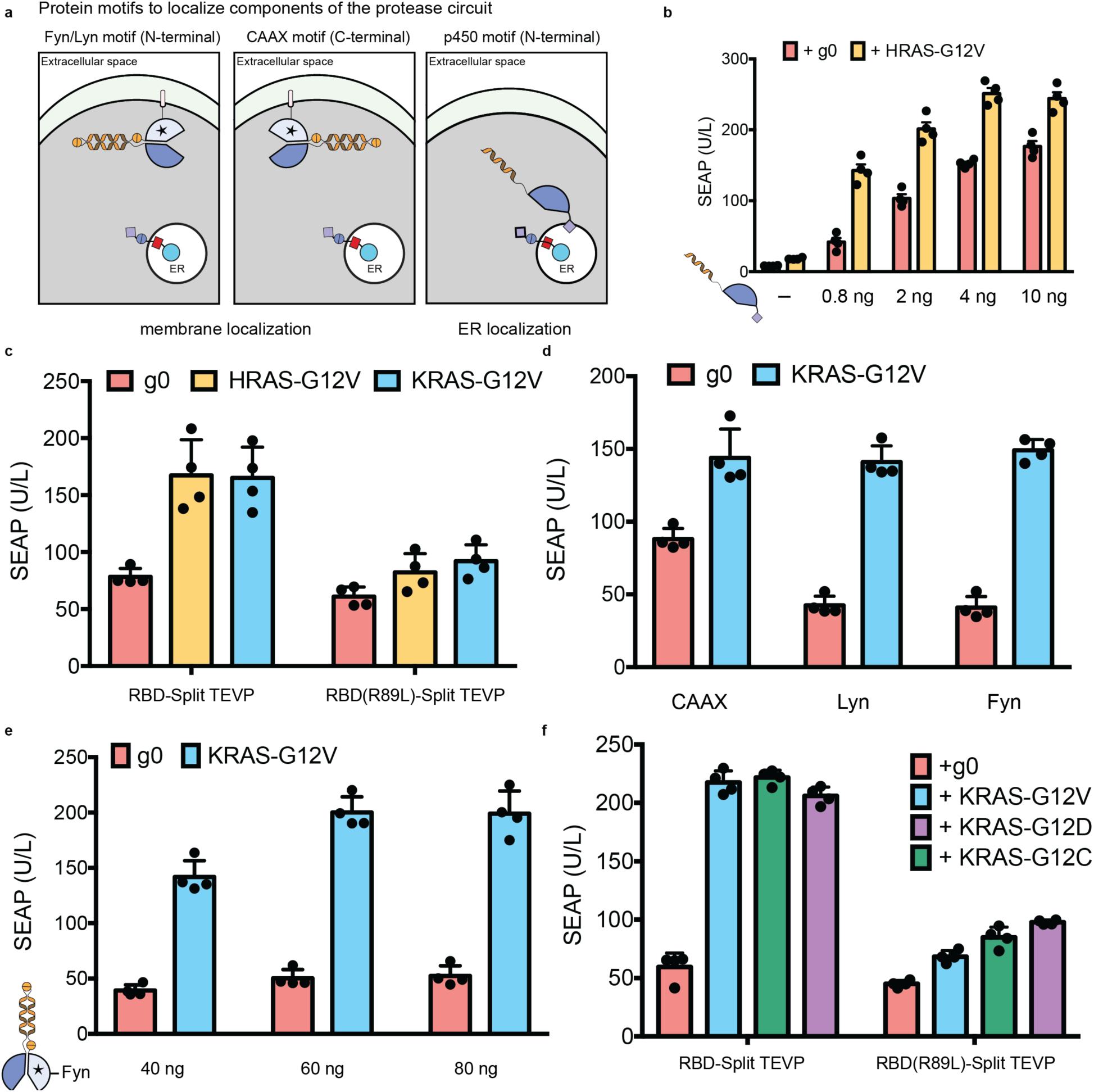
**a)** To efficiently propagate information from the cell membrane to the ER, signalling motifs were incorporated to localize components of the intermediate protease to the membrane via the Fyn/Lyn motif (left panel), or CAAX motif (middle panel). In addition, we used the signal anchor sequence of cytochrome p450 to localize components to the ER membrane (right panel). **b)** To increase the dynamic range of the RAS-sensing circuit (topology 3 from **Fig. 4d**), the amount of the ER-localized split TVMVP was reduced. **c)** The RAS-sensing circuit was comparable for sensing other RAS isoforms, such as KRAS-G12V. **d)** The Fyn and Lyn membrane associating motifs had reduced background relative to the CAAX motif, and **e)** increasing the amount of the membrane-associated TVMVP half localized with the Fyn motif, improved the dynamic range. **f)** The RAS-sensing circuit sensed other active mutants of KRAS at comparable levels to the KRAS-G12V mutant. Each dot represents a biological replicate. Mean values were calculated from four biological replicates (**b-f**) +/- SEM. The results are representative of at least two independent experiments; significance was tested using an unpaired two-tailed Student’s *t*-test between the two indicated conditions for each experiment. ** = p < 0.01, *** = p < 0.001, **** = p < 0.0001.

**Supplementary Table 1:** Apparent cleavage efficiencies of different RELEASE constructs used in this study. The apparently cleavage efficiencies of different RELEASE constructs were calculated by performing non-linear regression using the Michaelis-Menten equation. The cleavage efficiencies were represented by the K_m_ calculated from the fitted line. All non-linear regression was calculated using Prism 7.0.

| <u>RELEASE Plasmid</u> | <u>Protease Cut Site</u> | <u>Apparent Cleavage Efficiency</u><br><u>(amount (ng) of plasmid to</u><br><u>achieve $\frac{1}{2}$ Vmax) +/- S.E.</u> |
| --- | --- | --- |
| CMVTO-SEAP-26Sfur-3TM-tevs-KKMP | TEVP | 15.8 ± 1.6 |
| CMVTO-SEAP-26Sfur-B2AD-tevs-KKMP | TEVP | 1.1 ± 0.2 |
| CMVTO-SEAP-26Sfur-3TM-hcvs-KKMP | HCVP | 0.6 ± 0.1 |
| CMVTO-SEAP-26Sfur-B2AD-hcvs-KKMP | HCVP | 1.7 ± 0.2 |
| CMVTO-SEAP-26Sfur-3TM-hcvs-KKMP (GS) | HCVP | 5.5 ± 0.7 |
| CMVTO-SEAP-26Sfur-B2AD-hcvs-KKMP (GS) | HCVP | 113.9 ± 25.2 |
| CMVTO-SEAP-26Sfur-B2AD-tvmvs-KKMP | TVMVP | 7.7 ± 0.9 |

**Supplementary Table 2:**List of plasmids and the amounts use in this study. Please see attached excel file.

