## Supplemental Table 2 for "Protease-controlled secretion and display of intercellular signals"

| Figure 1 | Plate (wells) | ay length (h | Transfected plasmid (ng / well) | | | | | |
|---|---|---|---|---|---|---|---|---|
| | | | CMVTO-SEAP-26Sfur-3TM-tevs-AAMP | CMVTO-SEAP-26Sfur-3TM-tevs-KKMP | CMVTO-SEAP-26Sfur-B2AD-tevs-KKMP | CMVTO-SEAP-26Sfur-3TM-hcvs-KKMP | CMVTO-SEAP-26Sfur-3TM-hcvs-KKMP (GS) | CMVTO-SEAP-26Sfur-B2AD-hcvs-KKMP |
| 1c | 96 | 48 | 40 ng | 40 ng | / | / | / | / |
| 1d | 96 | 48 | / | 40 ng | / | 40 ng | / | / |
| 1e | 96 | 48 | / | 40 ng | 40 ng | / | / | / |
| 1f | 96 | 48 | / | / | / | 40 ng | 40 ng | 40 ng |
| 1h | 24 | 48 | / | / | / | / | / | / |

| Figure 2 | Plate (wells) | ay length (h | Transfected plasmid (ng / well) | | | | | |
|---|---|---|---|---|---|---|---|---|
| | | | CMVTO-SEAP-26Sfur-3TM-tevs-KKMP | CMVTO-GFP-26Sfur-3TM-hcvs-KKMP | CMVTO-SEAP-26sFur-B2AD-hcvs-tevs-KKMP | CMVTO-p450-hcvs-B2AD2-26Sfur-SEAP-26sFur-B2AD-tevs-KKMP | CMVTO-SEAP-26SFur-B2AD-tvmvs-KKMP | CMVTO-nTVMVP-AP4-tevs-P3-tevs-cTVMVPmut |
| 2a, b | 24 | 48 | 100 ng | 100 ng | / | / | / | / |
| 2c | 96 | 48 | / | / | 30 ng | / | / | / |
| 2d | 96 | 48 | / | / | / | 30 ng | / | / |
| 2e, f | 96 | 48 | / | / | / | / | 30 ng | 40 ng |

| Figure 3 | Plate (wells) | ay length (h | Transfected plasmid (ng / well) | | | | | |
|---|---|---|---|---|---|---|---|---|
| | | | CMVTO-IL12-26Sfur-B2AD-tvmvs-KKMP | TVMVP | CMVTO-mCherry-Kir2.1-HA-tevs-RXR | CMVTO-GFP-Kir2.1-HA-tevs-RXR | TEVP | pCAGGS-ASAP3 |
| 3a, b | 96 | 48 | 30 ng | 70 ng | / | / | / | / |
| 3c, d, e | 96 | 48 | / | / | 20 ng | / | 8 ng | 40 ng |
| 3f | 24 | 48 | / | / | / | 100 ng | 100 ng | / |

| Figure 4 | Plate (wells) | ay length (h | Transfected plasmid (ng / well) | | | | | |
|---|---|---|---|---|---|---|---|---|
| | | | CMVTO-SEAP-26SFur-B2AD-tvmvs-KKMP | CMVTO-nTVMVP-AP4-tevs-P3-tevs-cTVMVPmut | CMVTO-nTVMVPmut-tevs-AP4-tevs-P3-cTVMVP | CMVTO-nTVMVp-AP4-tevs-P3-tevs-cTVMVPmut-CAAX | CMVTO-p450-P3-cTVMVP | CMVTO-Fyn-nTVMVpmut-tevs-AP4-tevs-P3-cTVMVp |
| 4b, c | 96 | 48 | 30 ng | 20 ng | 20 ng | 40 ng | 0.8 ng | / |
| 4 d, e | 96 | 48 | 30 ng | / | / | / | / | 60 ng |

| Figure 5 | Plate (wells) | ay length (h | Transfected plasmid (ng / well) | | | | | |
|---|---|---|---|---|---|---|---|---|
| | | | CMVTO-SEAP-26SFur-B2AD-tvmvs-KKMP | CMVTO-p450-nTVMVp-AP4 | CMVTO-Fyn-nTVMVpmut-tevs-AP4-tevs-P3-cTVMVp | CMVTO-FKBP-GpA-nTEVp | CMVTO-FRB-CD28-cTEVp | CMVTO-IL12-26Sfur-B2AD-tvmvs-KKMP |
| 5b | 96 | 48 | 30 ng | 0.8 ng | 60 ng | 10 ng | 10 ng | / |
| 5c | 96 | 48 | / | 0.8 ng | 60 ng | / | / | 30 ng |
| 5d | 96 | 48 | / | 0.8 ng | 60 ng | 10 ng | 10 ng | 30 ng |
| 5e | 96 | 48 | / | 0.8 ng | 60 ng | 10 ng | 10 ng | / |
| 5f, g | 96 | 48 | 30 ng | / | 60 ng | / | / | / |

| Fig. S1 | Plate (wells) | ay length (h | Transfected plasmid (ng / well) | | | |
|---|---|---|---|---|---|---|
| | | | CMVTO-SEAP-26SFur-B2AD-tvmvs-KKMP | TVMVP | g0 | H2b-mCherry |
| S1 | 96 | 48 | 30 ng | 70 ng | 96 - 166 ng | 4 ng |

| Fig. S2 | Plate (wells) | ay length (h | Transfected plasmid (ng / well) | | | |
|---|---|---|---|---|---|---|
| | | | CMVTO-p450-hcvs-B2AD2-26Sfur-SEAP | HCVp | g0 | H2b-mCherry |
| S2 | 96 | 48 | 30 ng | 70 ng | 96 - 166 ng | 4 ng |

| Fig. S3 | Plate (wells) | ay length (h | Transfected plasmid (ng / well) | | | | | |
|---|---|---|---|---|---|---|---|---|
| | | | CMVTO-SEAP-26Sfur-B2AD-tevs-KKMP | CMVTO-SEAP-26Sfur-3TM-tevs-KKMP | CMVTO-p450-FKBP-nTEVp | CMVTO-p450-FRB-cTEVp | g0 | H2b-mCherry |
| S3b | 96 | 48 | 30 ng | / | 10 ng | 10 ng | 146 ng | 4 ng |
| S3c | 96 | 48 | / | 30 ng | 10 ng | 10 ng | 146 ng | 4 ng |

| Fig. S4 | Plate (wells) | ay length (h | Transfected plasmid (ng / well) | | | | | |
|---|---|---|---|---|---|---|---|---|
| | | | CMVTO-SEAP-26Sfur-B2AD-SpyTag-tvmvs-KKMP | CMVTO-p450-hcvs-SpyCatcher | HCVP | TVMVP | g0 | H2b-mCherry |
| S4 | 96 | 48 | 10 ng | 60 ng | 60 ng | 10 ng | 56 - 126 ng | 4 ng |

| Fig. S5 | Plate (wells) | ay length (h | Transfected plasmid (ng / well) | | | | |
|---|---|---|---|---|---|---|---|
| | | | CMVTO-mCherry-Kir2.1-HA-tevs-RXR | CMVTO-mCherry-Kir2.1-HA-tevs-KKMP | TEVP | g0 | CMVTO-BFP |
| S5 | 96 | 48 | 20 ng | 20 ng | 8 ng | 132 - 140 ng | 40 ng |

| Fig. S6 | Plate (wells) | ay length (h | Transfected plasmid (ng / well) | | | | | |
|---|---|---|---|---|---|---|---|---|
| | | | CMVTO-RBDK65E-nTEVp | CMVTO-RBDK65E-cTEVp | CMVTO-HRAS-G12V | CMVTO-SEAP-26Sfur-B2AD-tevs-KKMP | g0 | H2b-mCherry |
| S6a, b | 96 | 48 | 0 - 60 ng | 0 - 60 ng | 0 - 20 ng | 30 ng | 86 - 166 ng | 4 ng |

| Fig. S7 | Plate (wells) | ay length (h | Transfected plasmid (ng / well) | | | | | |
|---|---|---|---|---|---|---|---|---|
| | | | CMVTO-RBDK65E-nTEVp | CMVTO-RBDK65E-cTEVp | CMVTO-RBDK65E-R89L-nTEVp | CMVTO-RBDK65E-R89L-cTEVp | CMVTO-HRAS-G12V | CMVTO-KRAS-G12V |
| S7b | 96 | 48 | 10 ng | 10 ng | / | / | 0 - 20 ng | / |
| S7c | 96 | 48 | 0 - 10 ng | 0 - 10 ng | 0 - 10 ng | 0 - 10 ng | 0 - 20 ng | 0 - 20 ng |
| S7d | 96 | 48 | 10 ng | 10 ng | / | / | / | 0 - 20 ng |
| S7e | 96 | 48 | 10 ng | 10 ng | / | / | / | 0 - 20 ng |
| S7f | 96 | 48 | 0 - 10 ng | 0 - 10 ng | 0 - 10 ng | 0 - 10 ng | / | 0 - 20 ng |

Supplementary Table 2: List of plasmids and amounts used in study.

| CMVTO-SEAP -26Sfur-B2AD-hcvs-KKMP (GS) | CMVTO-GFP-ECL-3TM-tevs-KKMP V3 | CMVTO-GFP-ECL-3TM-hcvs-KKMP | TEVP | HCVP | g0 | H2b-mCherry |
|---|---|---|---|---|---|---|
| / | / | / | / | / | 156 | 4 ng |
| / | / | / | 0 - 60 ng | 0 - 60 ng | 96 - 156 ng | 4 ng |
| / | / | / | 0 - 60 ng | / | 96 - 156 ng | 4 ng |
| 40 ng | / | / | / | 0 - 60 ng | 96 - 156 ng | 4 ng |
| / | 50 ng | 50 ng | 150 ng | 150 ng | 290 ng | 10 ng |

| CMVTO-nTVMVPmut-tevs-AP4-tevs-P3-cTVMVP | TEVP | HCVP | g0 | H2b-mCherry |
|---|---|---|---|---|
| / | 0 - 125 ng | 0 - 125 ng | 25 - 275 ng | 25 ng |
| / | 0 - 60 ng | 0 - 60 ng | 106 ng | 4 ng |
| / | 0 - 60 ng | 0 - 60 ng | 106 ng | 4 ng |
| 40 ng | 0 - 30 ng | - | 50 - 86 ng | 4 ng |

| g0 | H2b-mCherry | CMVTO-BFP |
|---|---|---|
| 26 - 96 ng | 4 ng | / |
| 92 - 100 ng | / | 40 ng |
| 200 - 300 ng | / | 100 ng |

| CMVTO-p450-nTVMVp-AP4 | CMVTO-RBDK65E-nTEVp | CMVTO-RBDK65E-cTEVp | CMVTO-HRAS-G12V | CMVTO-KRAS | CMVTO-KRAS-G12V | CMVTO-RBDK65E-R89L-nTEVp | CMVTO-RBDK65E-R89L-cTEVp | g0 | H2b-mCherry |
|---|---|---|---|---|---|---|---|---|---|
| / | 10 ng | 10 ng | 0 - 20 ng | / | / | / | / | 86 - 106 ng | 4 ng |
| 0.8 ng | 10 ng | 10 ng | / | 0 - 20 ng | 0 - 20 ng | 10 ng | 10 ng | 65.2 - 85.2 ng | 4 ng |

| CMVTO-KRAS-G12V | CMVTO-RBDK65E-nTEVp | CMVTO-RBDK65E-cTEVp | CMVTO-p450-nTVMVp-hcvs-AP4 | CMVTO-p450-Ns4A-tv-HCVp-P3 | ASAP3 | CMVTO-HA-mCherry-Kir2.1-tevs-RXR | g0 | H2b-mCherry |
|---|---|---|---|---|---|---|---|---|
| / | / | / | / | / | / | / | 85.2 ng | 4 ng |
| 0 - 20 ng | 10 ng | 10 ng | / | / | / | / | 65.2 - 85.2 ng | 4 ng |
| / | / | / | / | / | / | / | 85.2 ng | 4 ng |
| / | / | / | / | / | 40 ng | 20 ng | 77.2 ng | / |
| 0 - 20 ng | 10 ng | 10 ng | 0.8 ng | 40 ng | / | / | 25.2 - 85.2 ng | 4 ng |

| CMVTO-KRAS-G12C | CMVTO-KRAS-G12D | MVTO-SEAP-26Sfur-B2AD-tvmvs-KKI | CMVTO-nTVMVp-AP4-tevs-P3-tevs-cTVMVPmut-CAAX | CMVTO-p450-P3-cTVMVP | -nTVMVpmut-tevs-AP4-tevs | Lyn-nTVMVpmut-tevs-AP4-tevs-P3- | CMVTO-p450-nTVMVp-AP4 | g0 | H2b-mCherry |
|---|---|---|---|---|---|---|---|---|---|
| / | / | 30 ng | 40 ng | 0.8 - 10 ng | / | / | / | 75.2 - 105.2 ng | 4 ng |
| / | / | 30 ng | 40 ng | 0.8 ng | / | / | / | 105.2 ng | 4 ng |
| / | / | 30 ng | 0 - 40 ng | 0 - 0.8 ng | 0 - 40 ng | 0 - 40 ng | 0 - 0.8 ng | 85.2 - 105.2 ng | 4 ng |
| / | / | 30 ng | / | / | 40 - 80 ng | / | 0.8 ng | 65.2 - 105.2 ng | 4 ng |
| 0 - 20 ng | 0 - 20 ng | 30 ng | / | / | 60 ng | / | 0.8 ng | 65.2 - 85.2 ng | 4 ng |
